# Challenging the classical view of CSF flow: measuring CSF net velocity in the human subarachnoid space with 7T MRI

**DOI:** 10.1101/2025.03.04.641359

**Authors:** E.C. van der Voort, M.C.E. van der Plas, M.W.J.M. Gosselink, J.J.M. Zwanenburg

## Abstract

Traditionally, cerebrospinal fluid (CSF) is believed to exit the brain via arachnoid villi, being absorbed into the superior sagittal sinus (SSS), with a net flow towards these exit sites driven by constant CSF turnover. However, measuring these velocities non-invasively in humans is challenging due to their slow nature and the presence of relatively large confounding factors such as physiological CSF pulsations (heartbeat and respiration) and head motion. This study presents a novel magnetic resonance imaging (MRI) method designed to measure the net velocity of CSF whilst accounting for confounding effects, which is called CSF displacement encoding with stimulated echoes (CSF-DENSE). By applying a similar model as used to study sea-level rise, different motion components of CSF were successfully disentangled. Simulations, along with phantom and in vivo experiments, demonstrate the ability of CSF-DENSE combined with time series analysis using unobserved components modeling to detect ultraslow velocities of approximately 1 μm/s, even in the presence of confounding motions that are an order of magnitude larger. If the major egress of CSF were via the SSS, the expected net velocity towards the SSS was estimated to be 4.22±0.14 µm/s, based on measured CSF net flow through the aqueduct into the subarachnoid space (SAS). However, no significant net velocity toward the SSS was observed (v = -0.18±0.15 µm/s, with positive velocity directed towards the SSS), thereby challenging the classical view of CSF outflow. These findings suggest the need to reconsider traditional models of CSF outflow pathways, with potential implications for understanding and treating neurological disorders.

## Introduction

The brain is a complex and highly metabolically active organ that requires efficient mechanisms to maintain brain homeostasis and guard its function. Cerebrospinal fluid (CSF) serves a crucial role in this process by facilitating the clearance of waste products and distributing signaling molecules such as hormones.[1] With the choroid plexus (CP) considered to be its primary excretion site,[2], [3] CSF circulates within the ventricular system and the subarachnoid space (SAS) and acts as a conduit for the transportation of waste products out of the brain parenchyma.[1] Understanding the dynamics of CSF and its role in brain clearance mechanisms is essential for elucidating the pathophysiology of neurological disorders. In recent years, substantial attention has been focused on understanding the driving forces behind CSF circulation. These driving forces, including cardiac pulsations, respiration, and low frequency oscillations (LFO), have been shown to affect the motion of CSF.[4], [5], [6], [7], [8], [9], [10] While these oscillatory motions arguably play a vital role in brain clearance,[8] the importance of CSF turnover remains underappreciated. CSF turnover, defined by Smets et al. as *“CSF production in relation to its distribution volume”*, is suggested to be a critical driver of brain clearance and is needed to refresh CSF in order to clear waste products.[11] The net velocity of CSF, induced by this continuous excretion, could be a target to assess the CSF turnover, provided that excretion and absorption occur at spatially distinct locations.

According to the classical view of CSF transport, CSF flows from its excretion site located at the lateral, third, and fourth ventricles via the foramina of Luschka and foramen of Magendie towards the SAS, where absorption into the superior sagittal sinus (SSS) occurs through the arachnoid villi (Figure 1).[1], [12], [13] However, it has been argued that this view is contradicted by various experimental studies and clinical data[14], and a more recent study has challenged this classical view, as exchange of water between blood and CSF was not only observed around the CP but also within cortical areas.[15] It was suggested that continuous water exchange may occur along the CSF pathways, implying that the CP may not be the primary site of CSF excretion.[14] Furthermore, anatomical evidence supporting the outflow of CSF through the arachnoid villi is very limited.[16] Specifically, studies show that the size, amount, and distribution of arachnoid villi change across the human lifespan[17] and their major development and maturation only occur after birth with full maturation at 18 months postnatal[18]. This suggests that, at least in early childhood, the arachnoid villi are not consistently used as the main exit route for CSF. Instead, alternative exit routes of CSF have been proposed, including pathways through the cribriform plate, along optic nerves and, via meningeal lymphatic vessels.[16], [19]

**Figure 1.**
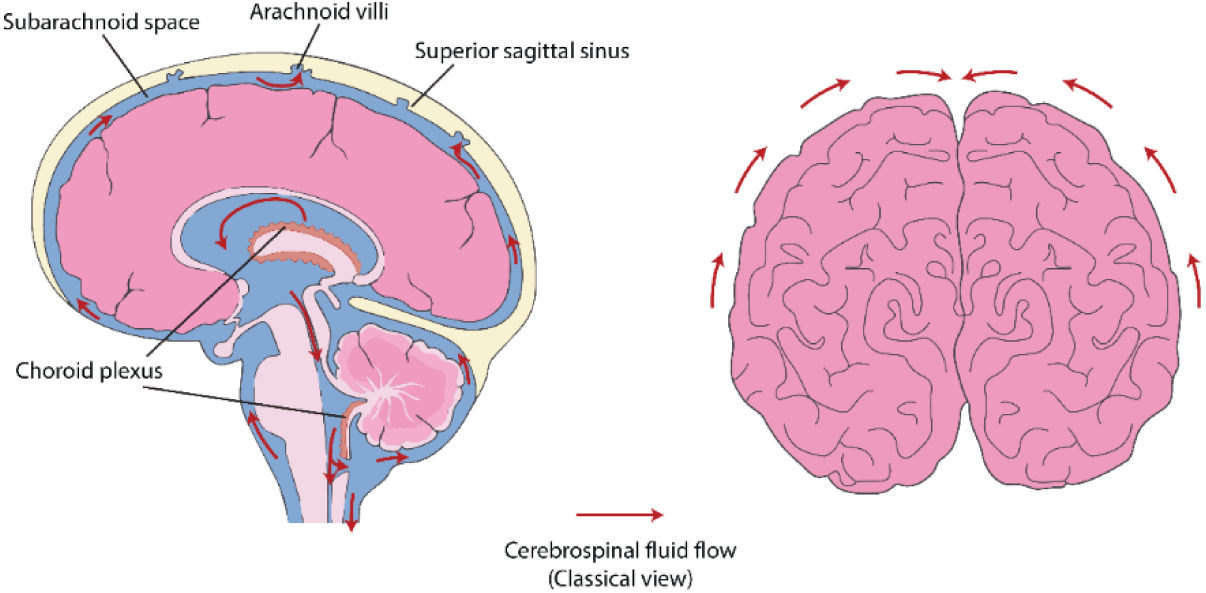
Schematic anatomical figure of the brain in sagittal (left) and coronal view (right). As described by the classical view of cerebrospinal fluid (CSF) flow, CSF is excreted by the choroid plexus and flows towards the subarachnoid space (SAS), where it is absorbed through the arachnoid villi into the superior sagittal sinus (SSS) (red arrows, left figure). The right figure shows the CSF flow in a coronal view, where just above the ears, CSF is expected to move in a Feet-Head (FH) direction in the SAS. At the top of the head the CSF is expected to move towards the SSS, therefore the flow is mainly in a Right-Left (RL) and vice versa (LR) direction (red arrows). This figure is adapted from Servier Medical Art, provided by Servier, licensed under a Creative Commons Attribution 4.0 unported license (https://creativecommons.org/licenses/by/4.0/).

While extensive rodent data has provided valuable insights into brain clearance mechanisms, translating these findings to humans remains challenging due to anatomical and physiological differences in CSF pathways and clearance.[1], [20] Magnetic resonance imaging (MRI) is the most suitable technique for acquiring data to measure and image CSF and brain clearance in humans since this is a non-invasive technique. However, current human in vivo data is limited and often relies on the use of contrast agents, of which correct interpretation can be challenging.[21] The ability to measure CSF net velocity due to CSF turnover would provide more insight into the CSF pathways, and potentially provide a tool to monitor CSF flows in disease. Previous studies measured the net CSF flow in cerebral aqueduct, which could serve as a proxy for the turnover of CSF by measuring the net inflow into the SAS.[22], [23] However, these measurements do not provide information on the absorption or possible exit routes of CSF. If CSF is indeed mainly absorbed via the arachnoid villi into the SSS, then the constant CSF turnover would result in a cranially and inwardly directed net velocity of CSF high in the SAS. A crude estimation of this net velocity based on the classical excretion rate by the CP and CSF volume shows that the net velocity of CSF would be around 5 μm/s in SAS for healthy young adults.[24]

Measuring these ultraslow velocities in vivo in humans is challenging. Magdoom et al. demonstrated that it is possible to measure flow velocities as slow as 1 µm/s using phase contrast (PC) MRI with stimulated echoes (STE) in a slow flow phantom.[25] They acknowledged, however, that in vivo measurements are more difficult due to periodic motions induced by the cardiac- and respiratory cycles. These periodic motions are estimated to be orders of magnitude larger than the net velocity at the base of the brain (close to the foramen magnum)[10], [26], [27] and induce CSF motions without contributing to a net displacement over time. Besides, head motion will be present which further challenges such measurements. Interestingly, the sequence used by Magdoom et al. is known in the field of cardiovascular MRI as Displacement Encoding with Stimulated Echoes (DENSE) and is used as a PC method for displacement mapping of the heart.[28] Conceivably, this sequence can be used to assess both the cardiac-related pulsatility and net velocity at the same time.[29], [30] Besides, research in the field of sea-level rise (approx. 2 mm/year or 5 µm/day) has shown that it is feasible to measure ultraslow velocities in the presence of large (tidal and seasonal) confounding variations in the measured data when sufficient data is available, and when the confounding factors are properly incorporated in the data analysis model.[31]

The aim of this study was to develop a method for measuring CSF net velocity resulting from CSF turnover using MRI, whilst simultaneously accounting for physiological induced motion such as heartbeat, respiration, and (restricted) head motion. Therefore, we combined an MRI method, called CSF Displacement encoding with stimulated echoes (CSF-DENSE), with a time series analysis using unobserved component modeling (UCM).[31], [32] First, to evaluate the performance of the developed method, simulations expanding on previous theoretical work were conducted.[24] Subsequently, measurements were performed in a motorized flow phantom containing small tubes with slow flow comparable to the expected CSF flow in the SAS. To further validate our method and show its feasibility in the presence of uncontrolled confounders, including head motion, additional phantom measurements were performed with a ground truth slow flow phantom attached to a human subject. Finally, as a proof of concept, in vivo data were acquired to estimate both the net velocity and periodic motions of CSF in the SAS. The focus of these in vivo measurements was on the upper regions of the SAS to determine whether a net velocity directed towards the SSS could be detected, to test the role of the SSS as a major outlet for CSF. We also compared the observed velocities with individual estimations of the expected velocity based on net flow estimations in the aqueduct as a proxy of the CSF excretion rate. Our results demonstrate that CSF-DENSE combined with UCM is capable of detecting very low velocities in the presence of serious confounding motions. The in vivo measurements challenge the classical view on the CSF outflow pathway via the arachnoid villi, as there was no significant net velocity of the CSF towards the SSS.

## Methods

### 1. Sequence Design

The proposed MRI sequence in this study represents a modification of the DENSE sequence previously used by Sloots *et al.* for the quantification of heartbeat-related brain tissue motion.(Sloots et al., 2020, 2021) The sequence comprises a non-selective motion encoding block and a slice selective motion decoding block. The time between the encoding and decoding blocks, known as the mixing time (TM), allows for the accumulation of the signal phase as a result of displacement of tissue and/or fluids (Figure 2). Importantly, during the TM, longitudinally stored magnetization decays with T1 (which is typically in the order of 1 second for biological tissues at 7T MRI). This is an essential difference compared to the widely used PC-MRI, where the motion-encoded signal decays towards the echo time (TE) with T2* (typically on the order of a few tens of milliseconds at 7T MRI). Therefore, T1 decay enables the use of relatively long TMs which are essential for sensitivity to slower flows. The motion sensitivity can be characterized by the displacement encoding parameter, DENC, which indicates the displacement that causes a phase shift of ±π.

**Figure 2.**
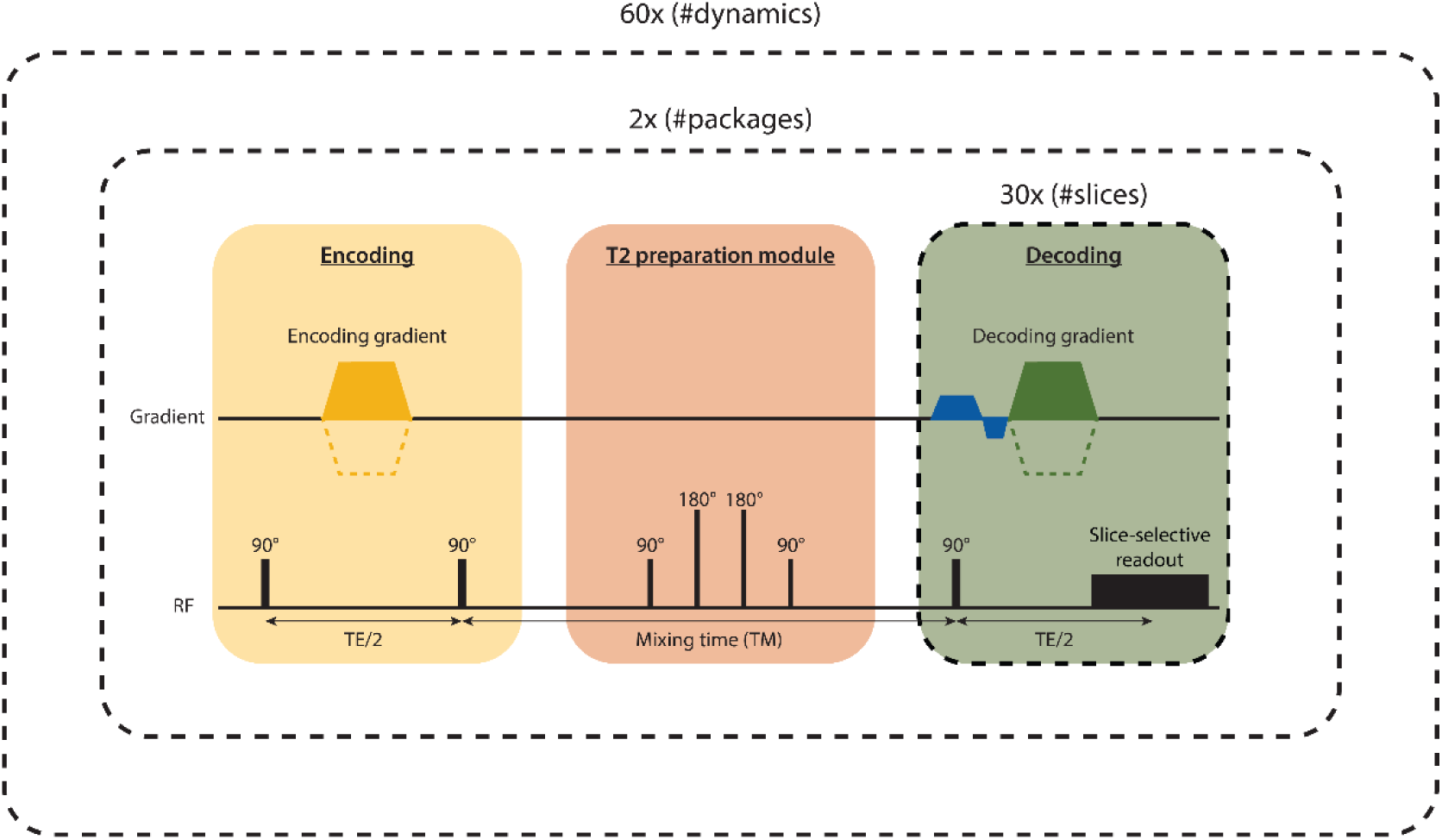
Schematic overview of the DENSE sequence. A non-selective encoding block (including two non-selective 90° RF pulses and the encoding gradient) is followed by a decoding block (including a slice-selective 90° RF pulse, decoding gradient with the same area as the encoding gradient and a readout). Phase accumulation measured during the mixing time (TM) corresponds to CSF displacement along the direction of motion encoding. In between encoding and decoding is a non-selective T2 preparation module which suppresses signals with short T2, to make the resulting signal specific for CSF. The imaging volume is split in two packages that contain the odd and even slices, respectively. The decoding block is repeated 30 times to cover the slices of one package in a single shot with increasing TM for every subsequent slice. The duration of one shot, consisting of encoding, T2 prep module and 30 slice decodings, is equal to one repetition time (TR). Two shots are needed to acquire both packages and to cover all 60 slices. In total, the sequence consists of 60 dynamics (repeats) in which the slices are systematically shuffled (so that every slice is acquired with the full range of TMs) and in which the gradient polarity is altered between the first 30 and last 30 dynamics (to allow for correction of eddy currents). The altered polarity is indicated by the dashed lines for the encoding/decoding gradients. The duration of the entire protocol is 12 minutes (2 x TR (6 sec) x 60 dynamics). TE = echo time.

Rather than focusing on tissue displacements, modifications were introduced to the previously used sequence to target CSF displacements. Firstly, adiabatic T2 preparation pulses (T2P) were introduced following the motion encoding block to render the signal CSF specific by utilizing the long T2 time of CSF[33] and thereby reducing partial volume effects. Secondly, the sequence was performed without triggering, using a fixed repetition time (TR), which yielded semi-random sampling of both the cardiac and respiratory contributions. This is needed to separate periodic physiological motions from a stationary motion component. Additionally, longer TMs were used to increase the sensitivity to phase shifts induced by the slow net CSF motion.

Accurately measuring the slow net velocity necessitates a combination of a high encoding sensitivity (low DENC) and a long TM to capture sufficient phase accumulation. However, a high encoding sensitivity and long TM reduce the SNR due to diffusion- and flow-induced signal loss.(Williamson et al., 2020) Furthermore, high sensitivities increase the risk of phase wraps resulting from cardiac and respiration-induced motions. Consequently, there is a balance between the chosen DENC and TM and their impact on the SNR. Based on preliminary in vivo assessments, a DENC of 0.125 mm provided a reasonable trade-off between the risk of phase wraps and sensitivity to motion, while a TM between 250 and 1900 ms was observed to provide a relatively high and sufficiently stable SNR. Besides, recently conducted simulations confirmed that such a relatively large range of TMs could accurately estimate different CSF motion components.[24]

For time efficiency, a single, non-selective encoding is followed by multiple slice-selective decodings (each decoding corresponds to the acquisition of one slice), where every decoding has an increasing TM. The sequence is repeated in a dynamic fashion, with slice shuffling implemented across the dynamics (repetitions) to ensure that every slice experiences every TM at least once. Following the completion of a cycle of slice shuffling, the dynamic series is repeated with altered polarity of the motion encoding and decoding gradients, in order to distinguish between motion-induced phase changes in the MRI signal and static phase confounders from e.g. the RF excitation pulses (Figure 2). The total number of dynamics should be enough to cover the full physiological cycle (i.e. each slice is measured multiple times at various moments in a given (cardiac or respiratory) cycle).[34]

The complete DENSE protocol acquires 60 slices, divided into two packages (odd and even slices). Each package has its own non-selective motion encoding block, resulting in 30 acquired slices (‘decodings’) per package. The slice interval was 60 ms, leading to TMs ranging from 250-1990 ms. The TR between the non-selective motion encoding blocks was 6s. A total of 60 dynamics were acquired (Figure 2): the first 30 with positive encoding and decoding gradient polarity, and the second 30 with negative polarity. The acquisition started with one dummy dynamic to reach steady state before the start of the data acquisition. With a TR of 6 seconds, the total scan duration is 12:12 minutes (6 s x 2 packages x (60 dynamics + 1 dummy dynamic)). The TE of the T2P module is 200 ms with a DENC of 0.125 mm and an acquired resolution of 3 mm isotropic. Detailed protocol information can be found in Table 1.

**Table 1.** MRI scan parameters.

|  | **T1 in vivo** | **T1 motorized flow phantom** | **DENSE in vivo** | **DENSE motorized flow phantom** |
| --- | --- | --- | --- | --- |
| **Orientation** | Coronal | Transversal | Coronal | Transversal |
| **Nr of slices** | 400 | 190 | 60 | 60 |
| **Nr of dynamics** | 1 | 1 | 60 | 60 |
| **Nr of packages** | 1 | 1 | 2 | 2 |
| **TMs (ms)** | - | - | 250 – 1990 | 250 – 1990 |
| **DENC (mm)** | - | - | 0.125 (FH/RL) | 0.25 (FH) |
| **T2 prep TE (ms)** | - | - | 200 | 50 |
| **EPI factor** | - | - | 33 | 35 |
| **SENSE factor** | 2 (AP); 2 (RL) | 2 (RL) | 2.6 (RL) | 2.3 (AP) |
| **Acquired resolution (mm<sup>3</sup>)** | 1.0 x 1.0 x 0.5 | 1.0 x 1.0 x 1.0 | 3.0 x 3.0 x 3.0 | 1.5 x 1.5 x 3.0 |
| **FOV (mm<sup>3</sup>)** | 200 x 252 x 200 | 250 x 250 x 190 | 250 x 250 x 180 | 120 x 120 x 180 |
| **Reconstructed in plane resolution (mm)** | 0.5 | 0.87 | 2.6 | 0.5 |
| **TR (ms)** | 9.0 | 4.2 | 6000 | 6000 |
| **TE (ms)** | 2.8 | 2.0 | 15 (TE/2) | 18 (TE/2) |
| **BW (Hz/pixel)** | 253.0 | 506.1 | 60.5 | 42.6 |
| **Total scan duration (min:s)** | 3:08 | 2:00 | 12:12 | 12:12 |

### 2. Simulations of the MRI protocol performance

To assess the performance of CSF-DENSE combined with UCM for distinguishing the different motion components, simulations were performed for a voxel in a single slice (see description of protocol above, and the parameters in Table 1). A fixed SNR of 15 (well below the minimum measured and achievable SNR with DENSE[29]) was used based on initial SNR assessments (data not shown). The measured phase data were generated from ground truth CSF displacement waveforms for the physiological processes (cardiac-, respiratory- and LFO-induced motion), for details of the ground truth generation and implementation of the simulation see[24].

To evaluate the CSF-DENSE combined with the UCM method in case of incorrect model assumptions, two simulations were performed with a mismatch between the underlying CSF dynamics and the assumed model. For the first simulation, the net velocity component occurred intermittently, with net flow occurring during the systolic phase of the cardiac cycle only (while a constant term over the full cycle was assumed in the analysis as described below). For the second simulation, LFOs were part of the CSF ground truth dynamics but not included in the model. The effect on net velocity estimation as well as the physiological processes (cardiac- and respiratory-induced motion) was evaluated. Both simulations included cardiac- and respiratory-induced motion. The first simulation included a net velocity of 10 μm/s restricted to the systolic phase, which corresponds to an effective net velocity of 4.64 μm/s over the full cardiac cycle (for an average systolic phase of 464 ms and an average cardiac cycle of 1000 ms). The second simulation included a continuous net velocity of 5 μm/s over the full cardiac cycle and LFO-induced motion. The amplitudes and cycle duration for the physiological processes were varied with amplitudes and durations of 100 ± 10 μm and 1000 ± 100 ms, 50 ± 15 μm and 5000 ± 1250 ms, and 75 ± 15 μm and 10 ± 2 s for the cardiac, respiratory and LFO cycle, respectively. A Monte Carlo approach was used to evaluate the noise-sensitivity of the estimated parameters with different noise realizations for each Monte Carlo run with a total of 1000 runs. The average and standard deviation (SD) of the estimated net velocity as well as the waveforms of the physiological processes was determined over all Monte Carlo runs.

### 3. Validation

#### 3.1 Motorized flow phantom

A 3D-printed phantom was designed to mimic both net flow and periodic motions (Figure 3). The phantom consisted of a cylinder filled with 2% agar gel containing two evenly spaced glass tubes (inner diameter = 4.0 mm, Wilmad, ATS) which were connected via a silicone tube outside of the gel. The end of one glass tube was connected to a syringe infusion pump (Fusion 100-X Syringe Pump, Chemyx) through an IV line, while the end of the other tube was placed in a container to collect the outflowing water. This cylinder was placed inside a sphere filled with a very concentrated solution of CuSO_4_ (which has an ultra-short T1 resulting in no detectable signal in the current protocol) which served to load the head coil and to provide some form of passive B0 shimming. The moving end of the phantom was attached to a motor (QUASAR™ MRI^4D^ Motion Phantom, Modus Medical Devices) located at the end of the scanner bed. The motor was strapped to the scanner bed to reduce (subtle) sliding during the periodic motion.

**Figure 3.**
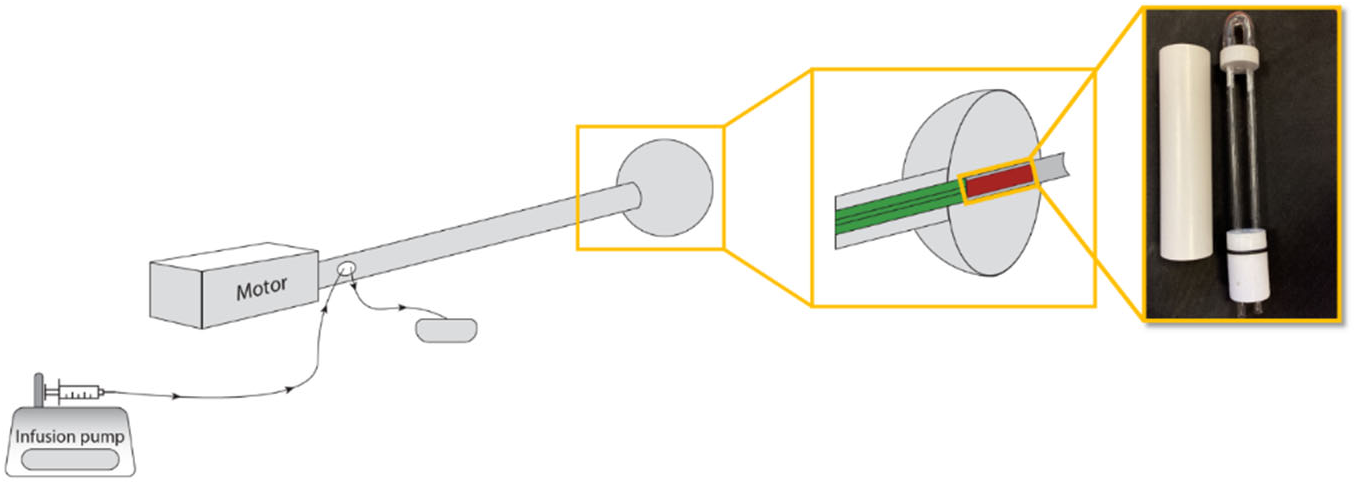
A schematic overview of the motorized flow phantom setup. The setup consists of a motor connected to a hollow pipe, which in turn is attached to a sphere. The first inset image shows the cross-sectional view of the sphere with the gel-filled cylinder (red) positioned inside the sphere, and the hollow, movable pipe (green). The second inset image shows the cylinder next to the two glass tubes for the transport of water. The glass tubes are inserted in this cylinder, which is subsequently filled with agar gel. Inside the hollow movable pipe (green) run two intravenous catheter lines, which are connected to the glass tubes. One catheter line is attached to a syringe and placed onto the infusion pump, the other is placed in a container to collect the outflowing water. The sphere is filled with a copper sulfate solution, which serves for loading of the head coil and provides passive B0 shimming. The motor is located at the end of the bed, with the sphere placed inside the head coil.

The infusion pump enabled a continuous water flow through the phantom. The rate of the pump was set to 0.05, 0.10, and 0.20 ml/h resulting in velocities of 1.11, 2.21, and 4.42 µm/s, respectively. The motor was able to move the cylinder horizontally along the Feet-Head (FH) axis (reference system aligned with a fictitious person being placed supine, head-first in the scanner). To mimic both cardiac and respiratory motion, two sinusoidal waveforms with different frequencies were applied to the motor. The cardiac frequency was set to 0.52 Hz and the respiratory frequency to 0.15 Hz. Both sinusoids were driven with the lowest possible nominal amplitude from the user interface of the motor, which was 0.1 mm.

Data were acquired using a 7T scanner (Philips Healthcare, Best, The Netherlands) with an 8-channel transmit and 32-channel receive head coil (Nova Medical, Wilmington, MA, USA). The DENSE data were acquired with motion encoding in the FH direction. All details of the MRI protocol can be found in Table 1. Physiological data traces were simulated using scanner-implemented software to match the settings of the motor, where the cardiac duration was set to 1935 ms (0.52 Hz) and the inspiration and expiration duration of the simulated respiratory signal were both set to 3333 ms, resulting in a frequency of 0.15 Hz. Additionally, a T1 scan was acquired for planning (see details in Table 1) and a T2* scan to detect the presence of potential air bubbles in the glass tubes with the following scan parameters: FOV = 120 x 120 x 120 mm^3^, resolution = 0.6 x 0.6 x 3 mm^3^, TR = 70 ms, TE = 18 ms, flip angle = 20°. Image-based shimming (second order) was performed on the agar gel of the cylinder.

#### 3.2 In vivo phantom

To validate the CSF-DENSE combined with the UCM method under similar conditions as in vivo, data were acquired in 2 healthy volunteers (2 females, 28.5 ± 0.7 years). The study was approved by the Ethical Review Board of our hospital, and written informed consent was obtained from each participant before scanning. The cylindrical part of the motorized flow phantom was strapped to the right side of the head of the volunteers to induce physiological motion in the phantom. The phantom was aligned in the FH direction, and foam padding was placed between the phantom and the head coil to avoid the transmission of mechanical vibrations of the scanner. A net flow was pumped through the phantom at a rate of 0.20 mL/h (corresponding to 4.4 μm/s flow velocity in the tubes). A transversal DENSE scan was acquired to assess the velocity in the phantom (see Table 1). For this scan, second-order image-based shimming was performed on the gel in the cylindrical phantom.

### 4 In vivo measurements

In vivo experiments were done in 6 healthy volunteers (5 males, 32.5 ± 8.1 years). The study was approved by the Ethical Review Board of our hospital, and written informed consent was obtained from each participant before scanning. DENSE data were acquired in coronal slices, with motion encoding in both FH as well as Right-Left (RL) direction. Detailed scan parameters can be found in Table 1. Physiological data were simultaneously acquired by using a peripheral pulse unit (PPU) based on a pulse oximeter, placed on the index finger, and a respiration belt, sampled at 500Hz. Volunteers were instructed to maintain a constant, calm abdominal breathing.

Additional acquired scans included a T1 scan for planning (Table 1 for scan parameters), 2D PC-MRI to determine the CSF net flow rate in the aqueduct (FOV = 190 x 250 mm^2^, resolution = 0.45 x 0.45 mm^2^, slice thickness = 3 mm, TR = 12 ms, TE = 4.5-5.9 ms (depending on angulation), flip angle = 12°, SENSE factor: 2 (Anterior-Posterior (AP)), VENC = 15 cm/s)[23] and a high resolution balanced steady-state free precession (SSFP) to determine the cross-sectional area of the SAS perpendicular to the FH encoding direction for calculating the expected net velocity (FOV = 250 x 240 x 200 mm^3^, resolution = 0.8 x 0.8 x 0.8 mm^3^, TR/TE = 5.0/2.5 ms, flip angle = 40°, SENSE factor: 2.3 (AP/RL)). Second-order image-based shimming was performed on the brain.

### 5 Data processing

#### 5.1 Preprocessing

Complex data were calculated from the phase data which were exported from the scanner. Several preprocessing steps were performed to correct for various error sources. An overview of the full data processing pipeline is presented in Figure 4. The pre-processing and analysis pipeline was implemented in MATLAB 2018b (The MathWorks, Inc., Natick, MA, USA).

**Figure 4.**
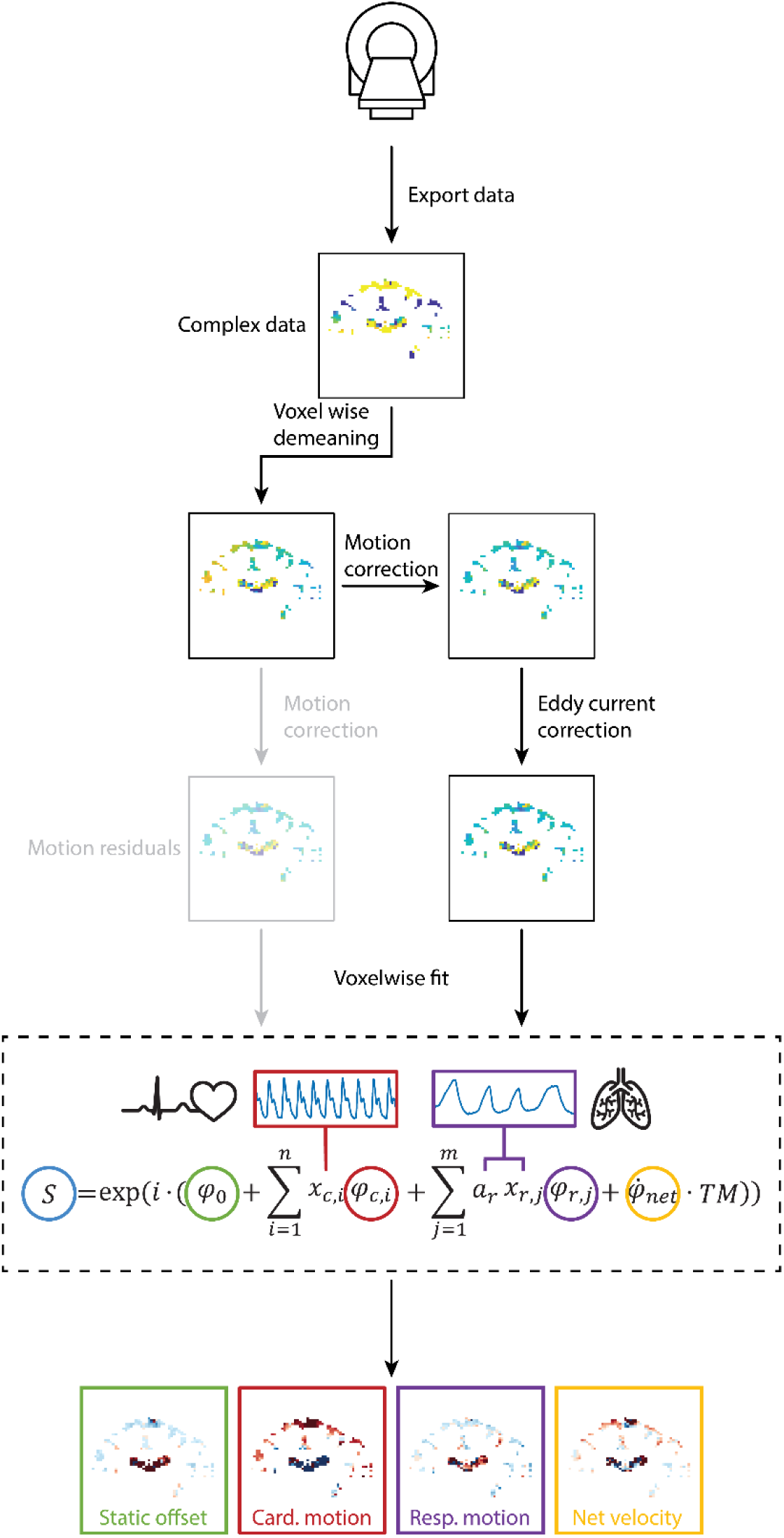
Overview of the processing pipeline. Data is retrieved from the scanner and the exported phase images are transformed to complex data. The first step is to temporally demean the data. Next, the data is corrected for rigid body motions due to involuntary head motion, and eddy currents. The final step is a voxelwise fit on the corrected data which yields a static offset map, cardiac and respiratory motion and net velocity map. The route in grey shows that the residuals of the motion correction can also be used as input for the voxelwise fit as the motion correction also partly removes physiological motions as far as these are present in the brain tissue (static tissue mask). For visualization purposes, the maps show the phase data, despite the fact that processing steps are applied to the complex data.

All corrections described below were calculated based on static masks and applied to both static and moving voxels. Moving masks (CSF/water) were made by selecting all voxels with a high signal intensity relative to the background, based on the magnitude images. Static masks (tissue/gel) were made by selecting all other voxels with an intensity above an empirically determined threshold (that selected voxels with an intensity above the noise level) and excluding the moving mask. For the in vivo data, the threshold for the moving mask was calculated by taking the 98% percentile of the maximum intensity. For the phantom data, the area of the tube corresponds to 50 voxels, given the spatial resolution. Therefore, per tube, the 50 voxels with the highest signal intensity were selected for the moving mask.

First, voxelwise demeaning of the phase over the 60 dynamics was performed to correct for the arbitrary, spatially varying phase offset that is always present in MRI phase data, thereby avoiding potential issues with phase wraps in the data. Demeaning was performed by taking the angle of the averaged complex data and adjusting the complex data for each slice and each dynamic accordingly. Note that averaging over all dynamics implies averaging data from both polarities of the motion encoding gradients, which, in principle, cancels out motion-induced phase variations. Still, as all motion in the series is measured with respect to a semi-arbitrary reference (i.e. the moment of motion encoding), adding or subtracting an arbitrary offset to the motion series does not affect the parameter fitting described in the next section.

Second, data were corrected for any rigid body motions such as rotation and translation induced by head motion that happened over TM (time between motion encoding and decoding). Rotational as well as translation movement results in a shift in the encoding direction according to:

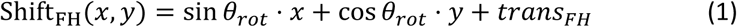

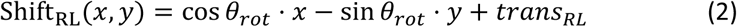

where *λ*_rot_ is the angle of rotation in degrees, *x*, *y* is the position of the voxel in mm and *trans_FH_*, *trans_RL_* is a translation along either the FH or RL direction in mm. The corresponding phase shift was determined according to:

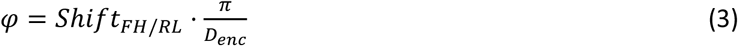

where *D_ene_* is the DENC of 0.125 mm for the in vivo data or 0.250 mm for the phantom measurements.

The rotation angle and translation were fitted using a nonlinear least-squares solver (*lsqnonlin*.m) with default settings and starting points equal to zero. This resulted in complex parameter estimates in which the real part represents the angle or translation and the imaginary part approaches zero. The angle and translation were estimated for each dynamic and every slice individually since the motion found in adjacent slices can be non-coherent due to the relatively high temporal spacing between slices caused by slice shuffling and the fact that odd and even slices are acquired in separate packages, one TR apart.

The in vivo data was only corrected for rotations along the through-plane axis (i.e. the AP axis) because data were acquired in coronal slices and thus the effect of rotations along other axes was much smaller (in a first-order approximation, rotations of a coronal plane around the RL and FH axes yield AP motions only). The phantom data were acquired in transversal slices with encoding along the FH axis. Since rotation around the through-plane axis (FH axis) does not result in motion along the encoding (FH) direction, the corrected shift was reduced to FH translations only.

Third, data were corrected for the presence of eddy currents (EC). The EC was estimated as an offset and a gradient in all three orthogonal directions and adjusted for voxel position with respect to the center of the magnet bore, according to:

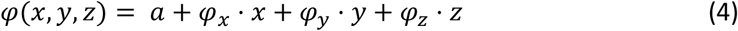

Where *a* is the offset in radians, *φ_x/y/z_* are the estimated EC gradients in every orthogonal direction in radians/mm and *x*, *y*, *z* are the 3D matrices with the voxel position in mm. The polarity of the encoding gradient was included in the voxel position matrices to account for positive and negative encoding polarities. All variables were estimated using the same nonlinear least-squares solver as before with a start value for the offset of 0.1 and for all EC gradients of 0.01.

#### 5.2 Parameter fitting

An adaption of a previously published model to estimate the sea-level rise in the presence of confounding effects from tides, atmospheric pressure, and wind was used to allow for the separation of cardiac and respiration-induced brain tissue motion and the small net velocity component.[31] In the current model, the total displacement is assumed to be a linear combination of the periodic motions due to the cardiac and respiratory cycles and net displacement due to CSF turnover. Therefore, the measured phase (*φ*) for a given voxel was modeled as follows:

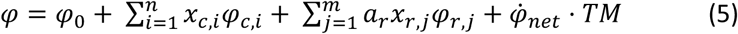

where *φ_O_* is a static phase offset, *φ_e_* and *φ_r_* are the phases due to cardiac and respiratory motion respectively, and *φ̇_net_* is the phase due to the net velocity. The *TM* represents the corresponding mixing time. Both cardiac and respiratory motion were retrospectively binned with *n* and *m* the amount of bins, respectively. A total of 10 bins was chosen for both periodic motions so that the data is estimated at 5, 15, 25,… and 95% of the full cycle. The binning is based on the physiological trace, where *x*_e_and *x*_r_ are the weights based on the temporal position of the measurements with respect to the cardiac or respiratory intervals, respectively. The additional weight, *a*_r_, is a weighting factor for the respiration cycle, based on the amplitude of the respiratory waveform during the interval in which *φ* was measured. This is under the assumption that the intracranial CSF flow scales with the depth of the respiration as seen by the respiratory belt.(Sloots et al., 2020) For simulations, this weight was set to 1, meaning that the variation in amplitude was not included in the model. Note that *φ_e_* and *φ_r_* are to be estimated per bin, without assuming a specific temporal waveform of the cardiac- and respiration-induced motions.

The different phase components were estimated by using a nonlinear least-squares solver with the normalized measured complex data in which the left side of equation (5) is replaced by its complex equivalent after applying all corrections, *e^iφ^*, and the right side is changed accordingly to avoid problems from potential phase wraps in the data. Default settings were used, with the following start values: 0.1 for the periodic motion, 0.01 for the net velocity component, and 1 for the static offset. The fit yields complex estimated parameters, of which the real parts were used as final results. Finally, the displacement (*y*) or velocity (*v*) can be derived from the estimated phase parameters by using the DENC (*D_enc_*):

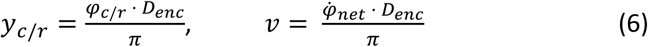

Since the data is acquired without triggering, the encoding and decoding moments fall at arbitrary positions within the cardiac and respiratory cycles. Therefore, the displacement is measured with respect to an arbitrary point in these cycles and a reference point was chosen by setting the first bin of both periodic motions to zero. Since, for the in vivo experiments, the PPU was used for measuring the cardiac cycle, the estimated cardiac curve was shifted by 3 bins (roughly 300 ms) for visualization purposes, in order to make the cycle start approximately at the ECG R-top (i.e. start of the systolic phase).

Applying the motion correction before EC corrections, improved the estimation of EC in the data. However, the motion correction removed the sinusoidal motion imposed by the motor on the phantom as well as the physiological motion in the phantom when placed next to a head, since the motion was equal in both the static gel and the flowing water in the tubes. To recover the periodic motion, the residuals of the motion correction step were fitted using equation (5) (see also Figure 4).

### 6 Data analysis

#### 6.1 ROI selection

For the phantom data (both motorized and in vivo), all slices with clearly visible glass tubes and homogenous B0-field were used for analysis, using the CSF mask as defined in section 5.1. Slices in which air bubbles were present, as visible on the T2* scan, were excluded. The mean and standard error of the mean (SEM) over all slices of the net velocity maps and of the physiological waveforms were determined.

The voxelwise fit yields a 3D net velocity map with isotropic 3 mm resolution voxels, that can be displayed and analyzed in any orientation. For in vivo data with encoding in the FH direction, the CSF net velocity was estimated in CSF voxels in the SAS located in 10 transversal slices, located just above the ears, where flow would be mainly in the FH direction (Figure 1), as these slices are approximately perpendicular to the skull and, thus, SAS. For the data with encoding in the RL direction, the net velocity was estimated in CSF voxels located in the upper 5 transversal slices of the SAS, where flow would be mainly in the RL direction towards the SSS. ROIs of the left and right hemisphere were defined by taking all voxels left or right of the interhemispheric fissure, respectively. The interhemispheric fissure was manually outlined on the DENSE magnitude data, on a coronal slice located in the middle of the AP axis of the brain and copied to all other slices. The ventricles were excluded for analysis by manually drawing ROIs. The mean and SD over all voxels within all slices of these ROIs were calculated for the net velocity estimations. The physiological motion was estimated in both ROIs for both encoding directions and the mean and SD of the waveforms over all voxels within all slices of these ROIs were reported.

For the RL encoded data, the centrally directed net velocity was identified by subtracting the average net velocity in the right hemisphere from the one in the left hemisphere, which should result in a velocity which is twice the expected net velocity (as leftward velocity is positive and rightwards velocity is negative in our data, by convention). To determine whether the measured net velocity is significantly different from zero, a t-test was performed for the mean net velocity in FH direction and the mean centrally directed net velocity of all subjects.

#### 6.2 Estimation of CSF excretion rate

The average velocity curves of the aqueduct were determined after applying background phase corrections to the PC-MRI data. The average velocity curve was multiplied by the aqueduct area to obtain an estimate of the volumetric flow rate Q (expressed in μL/s) of the CSF excreted by the CP in the lateral ventricles. For details of the analysis pipeline, we refer to a previous publication.[23] The amount of CSF voxels in the SAS was determined by applying a user-defined intensity threshold to the balanced SSFP data for every subject. The threshold was applied to transversal slices that covered the same region as used for the DENSE FH analysis and the ventricles were manually excluded. The areas of the resulting masks were averaged over the slices to obtain a single estimated effective SAS cross-sectional area per subject. Using the CSF excretion rate and the SAS cross-sectional area, the expected CSF net velocity was calculated according to:

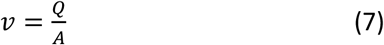

This velocity would be the expected net CSF velocity if all CSF excreted in the lateral ventricles would exit the cranium via the superior sagittal sinus as major outflow route.

## Results

### 1. Simulations of the MRI protocol performance

Simulations demonstrated that the CSF-DENSE protocol could accurately estimate the net velocity, even when there was a data-model mismatch. The estimated net velocity was 4.54 ± 5.28 μm/s which matched well with the ground truth (4.64 μm/s). The estimated cardiac and respiratory motions matched with the imposed ground truth curves (Figure 5) with a very slight underestimation. If LFOs are part of the total CSF dynamics but not taken into account in the model, the estimated net velocity was 5.14 ± 11.95 μm/s which matched with the ground truth of 5 μm/s albeit with a high SD. Figure 5 shows the estimated cardiac and respiratory curves which were slightly underestimated compared to the ground truth with relatively high SD.

**Figure 5.**
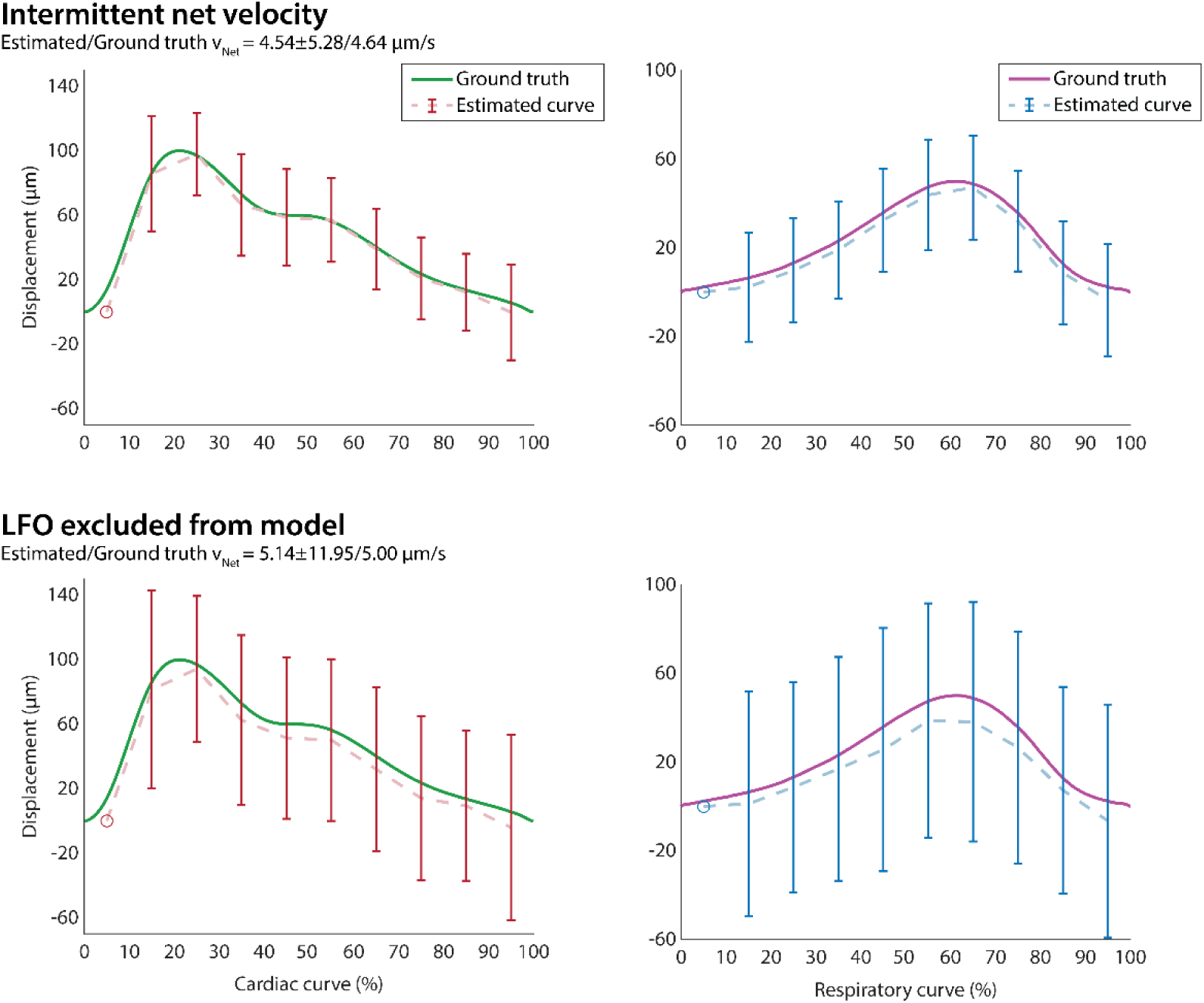
The estimated cardiac (left) and respiratory (right) waveforms and their ground truths for a simulation including cardiac- and respiratory-induced CSF motion with an intermittent net velocity (net flow limited to the systolic phase of the cardiac cycle only) (upper row) and for a simulation including cardiac-, respiratory- and LFO-induced CSF motion with constant net flow, but where the LFO was not included in the model (bottom row). The estimated waveforms were averaged over all Monte Carlo runs and the error bars indicate the SD over the Monte Carlo runs. Both waveforms were estimated with 10 bins which correspond to 5%, 15%, 25%, … and 95% of the ground truth where the first bin (5%), indicated by a circle, was used as a semi-arbitrary reference point which has zero displacement an no SD by definition. Both ground truths started at zero at 0% of the cycle, which explains the slight offset between the ground truth and the estimated waveforms.

### 2. Validation

#### 2.1 Motorized flow phantom

The B0 map was homogenous across 16 slices in the middle of the phantom. No air bubbles were observed in the T2* data in these slices, therefore all 16 slices were included for analysis. The average net velocities (mean ± SEM) for positive and negative flows at infusion rates 0.20, 0.10 and 0.05 mL/h were 4.84 ± 0.32 and -5.18 ± 0.15, 1.72 ± 0.18 and -1.75 ± 0.18 and 0.81 ± 0.17 and -1.29 ± 0.22 μm/s, respectively (Supplementary Figure 1).

Motion correction effectively removed the sinusoidal motion imposed by the motor and no motion was found after fitting the corrected complex data (Supplementary Figure 2). The two sinusoidal motion curves, retrieved after fitting the residuals of the motion correction, are shown in Figure 6 for all flow rates. The ‘respiratory’ motion curves (sinusoidal motion with frequency of 0.15 Hz) exhibited a range of 242.5 ± 2.8 µm, 164.9 ± 2.7 µm and 232.3 ± 3.5 µm (mean min-max difference ± SEM over all slices) at infusion rates of 0.20, 0.10 and 0.05 mL/h, respectively. The ‘cardiac’ motion curves, with a frequency of 0.52 Hz, consistently showed a smaller range than the ‘respiratory’ motion. The ranges for the waveforms were 79.4 ± 2.8 µm, 80.0 ± 2.8 µm and 89.5 ± 3.0 µm at infusion rates of 0.2, 0.1 and 0.05 mL/h, respectively. The start of each measurement was not synchronized with the motor. Therefore the initial phase of the motion waveforms varied across the different measurements as can be seen in Figure 6.

**Figure 6.**
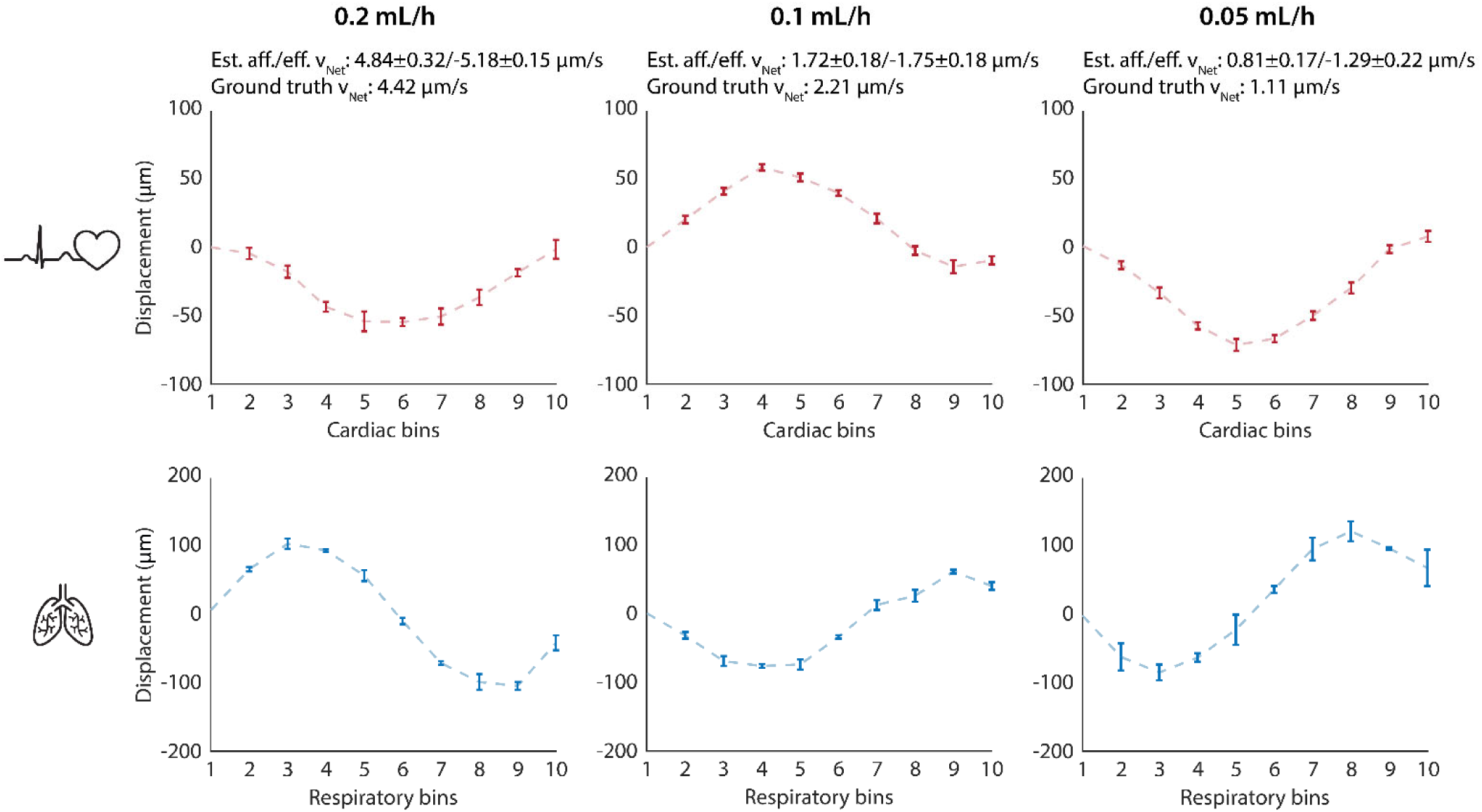
Results of motorized flow phantom experiment showing the physiological motion for both the cardiac (top row) and respiratory motion (bottom row). For different infusion rates: 0.20 mL/h (first column), 0.10 mL/h (middle column) and 0.05 mL/h (last column). The estimated net velocity for the afferent (Est. aff. v_Net_) (positive) and the efferent (Est. eff. v_Net_) (negative) tubes as well as the corresponding ground truth net velocities are depicted above the cardiac motion curves. Both motions were binned into 10 bins. The error bars indicate the SEM over all (n = 16) slices. The first bin is used as a reference point which has zero displacement and no SD by definition.

#### 2.2 In vivo phantom

Two subjects underwent scanning with the phantom placed adjacent to their heads. For both subjects, a gradient was observed in the net velocity maps of the phantom as shown in Figure 7A. The same gradient was found in isolated phantom measurements when the phantom was placed at an incline (Supplementary Figure 3). The measured net velocity in the phantom for these two subjects were 2.40 ± 0.16, -2.92 ± 0.17 μm/s, and 1.30 ± 0.26, -4.25 ± 0.29 μm/s in the afferent and efferent tubes of the phantom, respectively (Figure 7B). The cardiac and respiratory waveforms of the phantom are shown in Figure 7C. The cardiac curve showed a typical cardiac waveform for subject 1 and the respiratory curves revealed that the phantom moves downwards with every inhalation.

**Figure 7.**
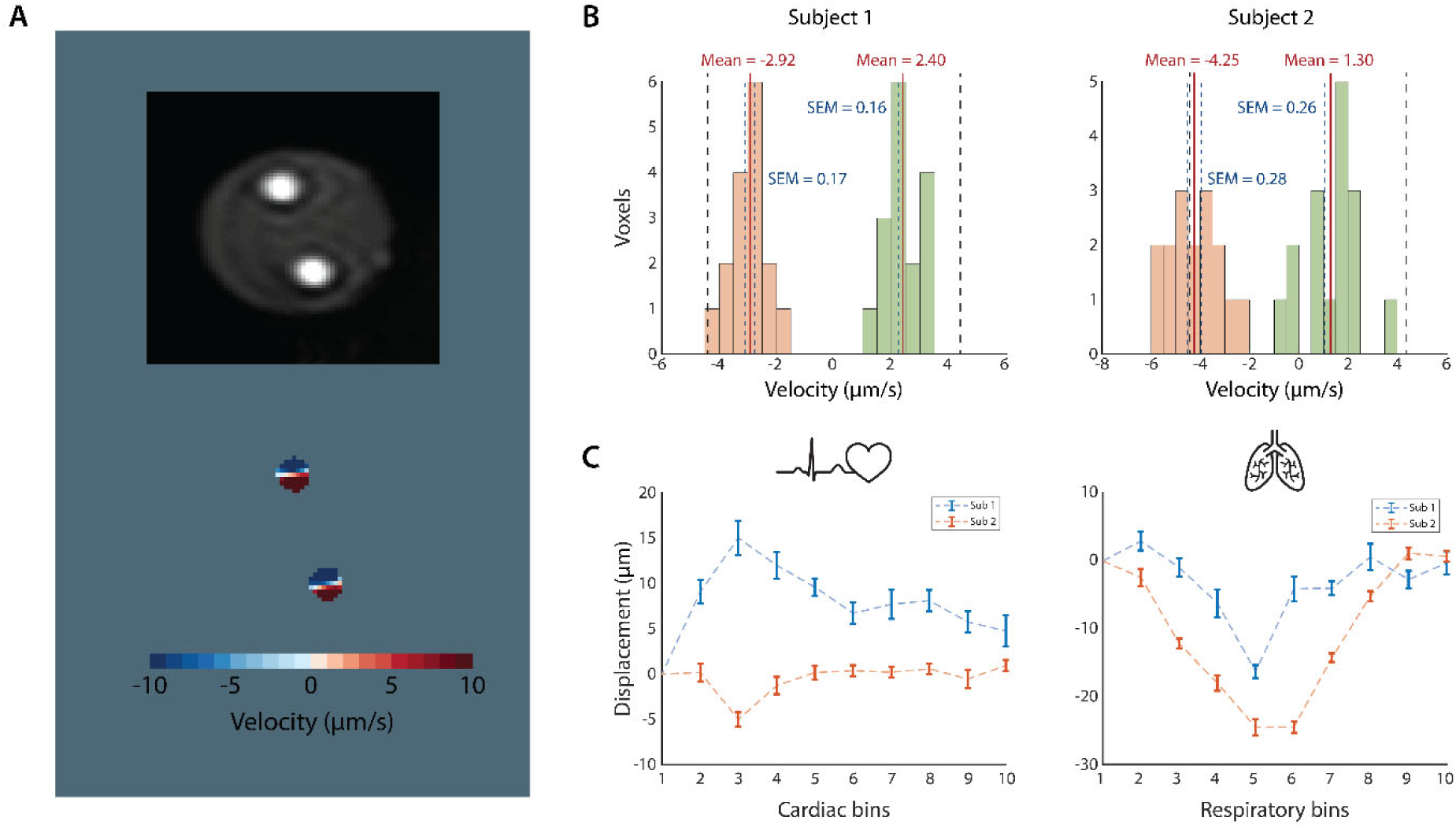
In vivo phantom validation FH-velocity results. A) Example of the gradient visible in the net FH-velocity maps from subject 1. This gradient was found in most slices in both subjects and could be reproduced in isolated phantom experiments when the phantom was placed at an incline (see Supplementary Figure 3), suggesting that vibrations in combination with gravitational forces induced a vortex. B) Histograms of the mean velocity of all slices for subjects 1 (left) and 2 (right). The red line indicates the mean velocity, the dashed blue line the SEM over all slices and the black dashed line the ground truth velocity. The orange and green histograms represent the voxels in the efferent and afferent tubes, respectively. C) The averaged cardiac (left) and respiratory (right) motion in the residuals after motion correction over all slices for both subjects. The error bars indicate the SEM over all (n = 16) slices. The phantom in subject 1 was placed higher and with a different orientation compared to subject 2, possibly explaining the difference in direction sign of the cardiac motion.

### 3. In vivo measurements

The net velocity maps for both encoding directions for all subjects are presented in Figure 8. Table 2 shows the average net velocities (Histograms of all in vivo results are shown in Supplementary Figure 4). The measured net velocities in the FH direction were not significantly different from zero (p = 0.26, Table 2). In both the FH and RL encoding direction, the estimated net velocities did not exhibit a discernible directional pattern consistent with CSF flow towards the SSS, i.e. no upward or inward net CSF motion was found (Figure 8). No significant inwardly directed velocity was observed in the RL velocity maps (p = 0.71, Table 2 and Supplementary Table 1). The measured flow in the aqueduct is listed in Table 2. Based on these flow measurements and assuming the SSS as major outlet, a net velocity of CSF in the FH direction in the SAS of 4.22 ± 0.14 μm/s (mean ± SEM) was to be expected.

**Figure 8.**
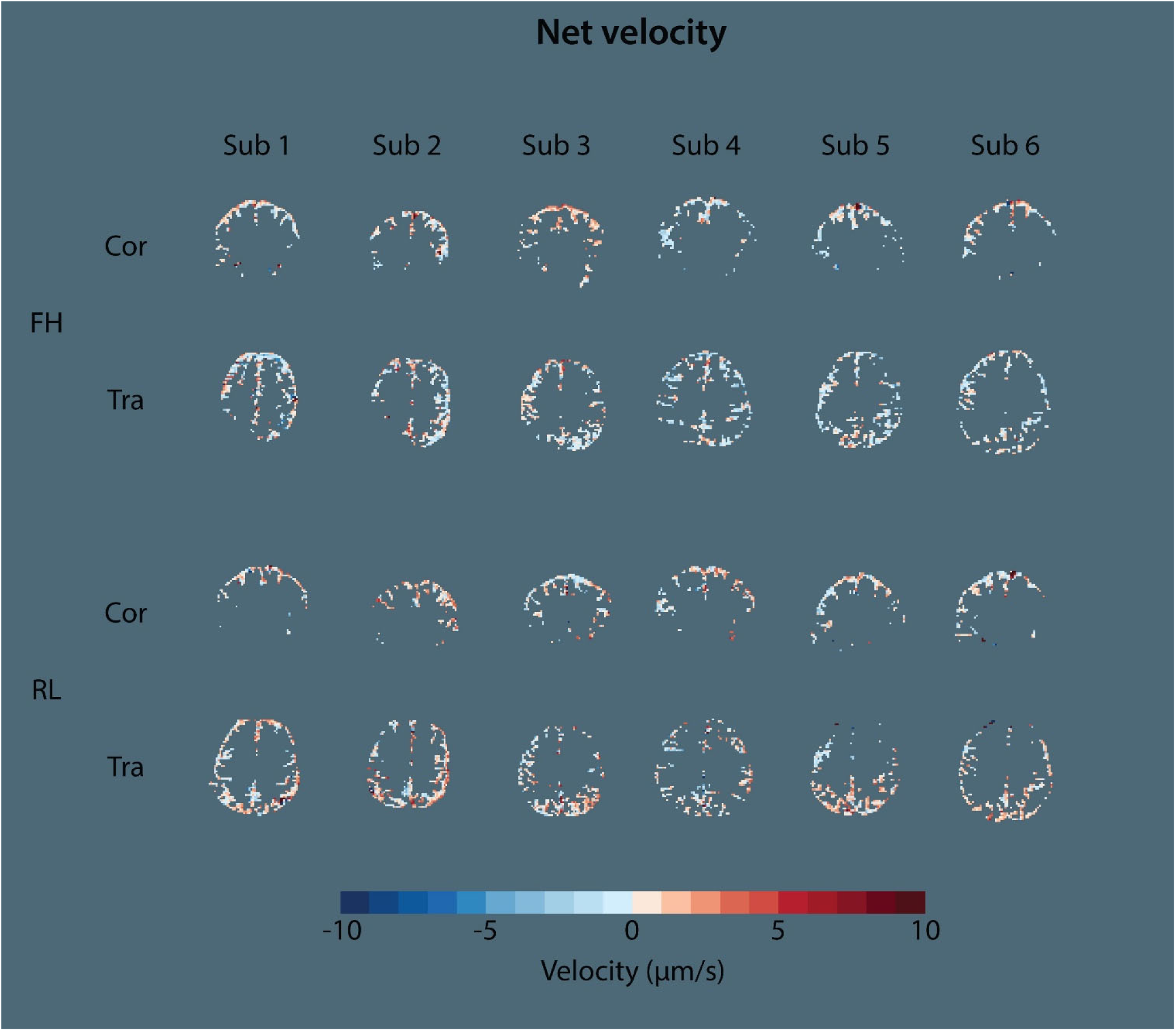
Overview of the voxelwise estimated net velocity results in vivo. Coronal (Cor) and transversal reformatted (Tra) slices of the estimated 3D net velocity maps for both feet-head (FH) and right-left (RL) velocities for all subjects. Positive velocities indicate dorsal or leftward flow for the velocity maps obtained with FH and RL encoding, respectively. See Supplementary Figure 4 for histograms and mean velocities for all subjects.

**Table 2:** Measured net CSF velocities in the SAS and CSF net flow through the aqueduct.

| | Measured FH net velocity ($\mu\text{m/s}$) | Measured centrally directed velocity ($\mu\text{m/s}$) | SAS cross-sectional area ($\text{mm}^2$)* | Flow aqueduct ($\mu\text{L/s}$)** | Expected FH net velocity in SAS ($\mu\text{m/s}$)*** |
| --- | --- | --- | --- | --- | --- |
| **Sub 1** | -0.6 | -0.7 | 2058 | -8.0 | 3.9 |
| **Sub 2** | -0.4 | 0.0 | 2100 | -10.0 | 4.8 |
| **Sub 3** | 0.4 | 0.1 | 1455 | -5.7 | 3.9 |
| **Sub 4** | -0.4 | -0.3 | 2579 | -11.0 | 4.3 |
| **Sub 5** | 0.0 | 1.0 | 1818 | -7.7 | 4.2 |
| **Sub 6** | -0.1 | 0.4 | 1791 | -7.6 | 4.2 |
| **Mean ± SEM** | $-0.18 \pm 0.15$ | $0.08 \pm 0.24$ | $1967 \pm 154$ | $-8.33 \pm 0.77$ | $4.22 \pm 0.14$ |

\*Estimated from balanced SSFP scans
\*\*Estimated from PC-MRI through the aqueduct
\*\*\*Computed using Equation 7

The physiological motion in the FH direction measured in the upper slices of the SAS and in the RL direction measured in the middle slices of the SAS are shown in Figure 9. The estimated cardiac motion exhibits a typical shape over the cardiac cycle, with CSF moving upward (FH) and inward (RL) during the systolic phase. The estimated respiratory curves indicated no clear CSF motion in either the FH or the RL direction during inspiration or expiration. Analysis of the residuals after correcting for head motion showed that in some cases, these corrections removed respiratory motion, as the residuals showed a ventrally directed CSF motion with respiration (Supplementary Figure 5).

**Figure 9.**
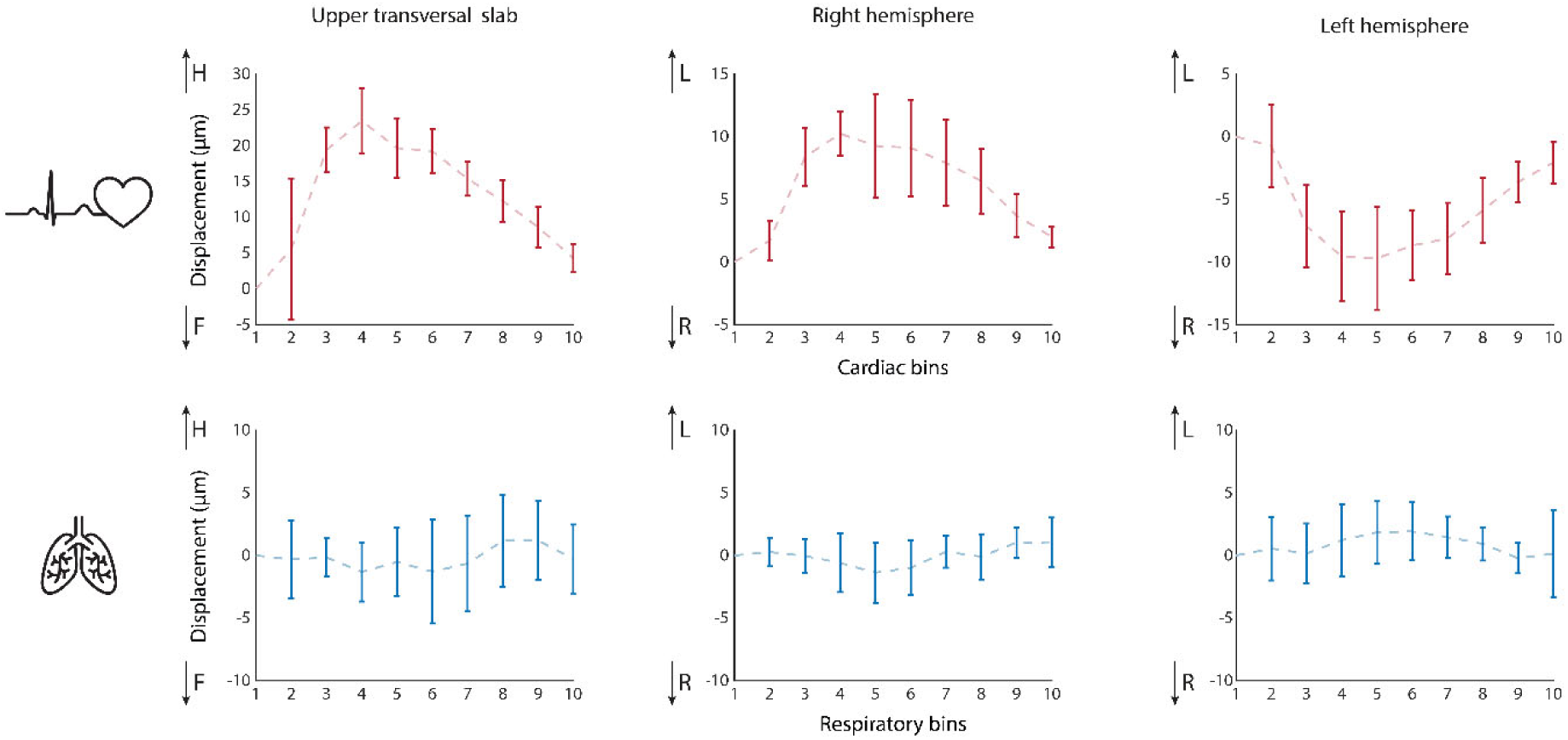
Estimated cardiac (upper row) and respiratory (bottom row) motion analyzed in the 5 upper transversal slices of the SAS for FH encoding (first column) or 10 transversal slices just above the ears in the right and left hemisphere for the RL encoding (second and third column, respectively). A typical cardiac curve is seen in CSF motion over the cardiac cycle where CSF moves cranially (FH) and inwardly (right/left) during the systolic phase. No clear CSF motion was measured for FH direction or between the right and left hemisphere over the respiratory cycle. Motion is binned into 10 bins which represent 5%, 15%, 25%, … up to 95% of the cardiac or respiratory cycle, bins 1-5 represent inhalation and bins 6-10 exhalation. The error bars indicate the SD over all (n = 6) subjects. The first bin is used as a reference point which has zero displacement and no SD by definition.

## Discussion

This study presents CSF-DENSE, a non-invasive method based on MRI, combined with time series analysis using UCM to measure CSF net velocity in the SAS in humans while accounting for head motion and periodic motion induced by heartbeat and respiration. The performance of CSF-DENSE combined with UCM was evaluated using simulations and further validation was obtained by measuring net velocities in a motorized flow phantom and a flow phantom under in vivo conditions. As a proof of concept, we focused the in vivo measurements on the upper regions of the SAS, because outflow is traditionally expected to occur via the arachnoid villi that protrude into the SSS[1], [12], [13]. Phantom measurements showed the ability of CSF-DENSE combined with UCM to detect net velocities as low as approximately 1 μm/s in the presence of periodic motions that were at least one order of magnitude larger. The in vivo results indicate no detectable net velocity towards the SSS (i.e. the averaged net velocity observed in the individual subjects was at most one-tenth of the velocity that was expected based on quantitative flow measurements of the net flow of CSF through the aqueduct into the SAS). These findings challenge the classical view of CSF absorption. This illustrates the potential of our method to measure CSF flow dynamics and to provide valuable insights into brain clearance mechanisms by helping to identify potential outflow routes of CSF, which could serve as a proxy for brain clearance.

### Related work

Several in vivo techniques currently exist that are able to non-invasively measure CSF dynamics, such as diffusion-weighted imaging[35], arterial spin labeling[15], real-time velocimetry[36], BOLD[6] and balanced SSFP[37]. While these methods provide valuable insights into the CSF dynamics, they do not target the net CSF flows that result from continuous CSF turnover. CSF-DENSE could be used as a direct, quantitative measure of CSF turnover. Existing MRI methods that measure net CSF flow through the aqueduct appear to provide reasonable estimates of the CSF excreted by the CP.[23], [38], [39], [40] However, they do not provide insight into the subsequent fate of the CSF that flows via the aqueduct into the SAS and are unable to address excretion at any other sites.

Previous studies have demonstrated the feasibility of measuring ultraslow net velocities in static phantoms[25], [41]. Our findings show that CSF-DENSE extends this capability by achieving comparable results in the presence of large confounding motions. These confounding motions (i.e. heartbeat and respiration) were correctly estimated in a phantom setup. However, our in vivo measurements showed no CSF motion due to respiration. Studies that did observe the respiratory-induced motion of CSF did so in very low regions of the brain, e.g. the aqueduct or fourth ventricle[23], [27], where the brain is known to pulsate much more than in the upper parts[42]. In higher regions, the CSF pulsations over the respiratory cycle were found to be lower[26], possibly explaining the absence of respiration-related CSF motions in our results.

### Validation

Recently, we demonstrated using simulations that the proposed approach is feasible. However, the implementation of CSF-DENSE in practical settings is not trivial. Nevertheless, the current work shows that the approach is both practically feasible and sufficiently sensitive to estimate very low net velocities in the presence of relatively large confounders. Analyzing time series with UCM has previously been utilized for estimating the rise in sea-level.[31] This successful application in a different field supports the reliability of the approach in our study, as it addresses analogous challenges, albeit within the context of CSF dynamics rather than oceanographic processes.

The motorized flow phantom results showed that CSF-DENSE is able to accurately measure slow net velocities in a controlled setup. To further validate the method, the phantom study was repeated in the presence of human subjects to account for ‘in vivo conditions’ such as motion, variability in heart rate and respiration, and B0 distortions. This demonstrated that even under these conditions, a significant net velocity was detected. However, a gradient was visible in the net velocity maps, likely due to the positioning of the phantom. The imperfect horizontal alignment of the phantom in combination with vibrations due to scanner noise may have allowed gravity to affect the flowing water, leading to vortex formation. When the experiment was repeated in isolation (without a human subject next to it) while the phantom was positioned at an incline, similar gradients were observed in the net velocity maps. The presence of these gradients probably impacted the accuracy of the net velocities estimations. Despite this gradient and the in vivo conditions, significantly different net velocities were measured in the afferent and efferent tubes of the phantom, and the average net velocities were much higher than those observed in the SAS. The net velocities in the individual voxels in the tubes were even much higher than typical velocities seen in humans, confirming that if any significant net velocity would be present, it should be discernable even on a voxelwise level. The presence of arachnoid trabeculae in the SAS may temper the dynamics of the CSF flow in the SAS as simulations suggested a much higher pressure drop with flow when trabeculae were incorporated in the computational fluid dynamics model.[43] This could explain the absence of similar scale vortices in the human SAS as were apparently present in the tubes of the flow phantom.

### Experimental challenges

Although the measured net velocity closely matched the ground truth value, the estimated amplitude of the high-frequency sinusoidal curve was lower than the nominal motor settings. This discrepancy is most probably due to the motor’s motion precision, which the vendor reported to be 0.25 mm (compared to the set amplitude of 0.1 mm). We observed that the motor did not reach the full amplitude, particularly at higher frequencies, probably as a result of slippage of the motor with respect to its surface. Therefore, lower frequencies were chosen in order to create robust confounders albeit at frequencies that were lower than the physiological cardiac and respiratory frequencies. Nonetheless, given that lower frequencies pose more challenges in distinguishing a net velocity (0 Hz) from periodic motions than high frequencies, these phantom measurements clearly demonstrated the feasibility of CSF-DENSE combined with UCM to measure very small net velocities in the presence of much larger confounders.

To estimate the expected net velocity in the SAS, the CSF flow in the aqueduct was measured as a proxy for the total CSF turnover. Flow measurements in the aqueduct likely underestimate the actual total CSF excretion because the excretion by CP located in the fourth ventricle is not included in these measurements. This would mean that the net velocity in the SAS should potentially even be higher if all excreted CSF would exit the SAS via the arachnoid villi in the SSS.

It is important to stress that the current sequence is highly sensitive to subtle motions in order to measure slow flows. This sensitivity also means, however, that any head motion has the potential to significantly affect the results. No macroscopic head motion was visible over dynamics and between slices hence registration between dynamics was deemed unnecessary. Nonetheless, correcting for subvoxel motion remained necessary. Motion correction is estimated based on tissue voxels, which still gave signal due to imperfect T2 preparation pulses. The motion correction was applied to the CSF voxels, assuming that the tissue voxels are static, i.e. have no net velocity, and that any interstitial fluid present in the tissue does not have a coherent, significant net velocity. Therefore, the measured physiological CSF motion represents motion relative to the brain tissue. This resulted in no visible respiratory motion of CSF, indicating that CSF does not move with the respiratory cycle relative to brain tissue. The residuals of the motion correction showed that, for some subjects, respiratory motion is filtered out due to the motion correction, similar to how motion correction filtered out the sinusoidal motion of the motorized flow phantom. The respiratory motion retrieved from the residuals showed CSF (and brain tissue) move ventrally (towards the feet) with every inhalation, consistent with previous brain tissue measurements that showed a rotation of the head with respiration causing the frontal region to move in the feet direction.[42]

### Implications

According to the classical view of CSF transport, there should be a net velocity in the SAS directed towards the SSS. Measurements of CSF flow in the aqueduct suggested that this velocity should be at least 4 μm/s. However, despite the fact that CSF-DENSE has been validated to be able to reliably measure these ultra-slow velocities, in vivo results revealed no detectable net velocity in the SAS, and certainly not close to the expected value.

Recently, more attention has been given to dual outflow where part of the CSF exits through venous routes and part through lymphatic routes.[44] This study demonstrates that there is no (or only very limited) net CSF absorption into the SSS. Therefore, it is likely that there must be another dominant exit route. It has been suggested that CSF can also exit the brain along cranial nerves, including the cribriform plate, or via meningeal vessels.[16], [44] Alternatively, inflow of CSF into brain tissue has been suggested by the glymphatic clearance theory.[45], [46] However, this would be accompanied by a gradient of decreasing net velocity from the base towards the top of the brain, which was not observed in the analyzed region of the SAS. The analyzed region is approximately halfway the SSS, which would imply that the velocity at this region would still be around 2 μm/s, which should be well detectable given the sensitivity and accuracy of our method. Another explanation for the absence of a net velocity in the SAS is the presence of a more concentrated net flow in the perivascular spaces surrounding blood vessels in the SAS.[47] However, the measured velocity in those perivascular spaces was in the order of 1 mm/min (roughly 20 μm/s). Considering the much smaller cross-sectional area of these perivascular spaces compared to the SAS, this velocity is still most likely too slow to accommodate all CSF excreted by the CP to move towards the SSS via this pathway. Hence, another exit route for CSF would be needed. At the same time, if alternative outflow routes exist in parallel to outflow via the SSS, it cannot be excluded that lower, but physiologically relevant net velocities in the SAS are present that are below the detection threshold of the current method.

In this work, we primarily focus on measuring the net velocity of CSF and use this to validate the classical hypothesis of CSF transport towards the SSS. Nevertheless, the current method is also capable of estimating the cardiac- and respiration-induced CSF motions,[24] which may play an important role in clearance mechanisms. While this approach targets the movement of solvent rather than the solutes, and therefore does not provide insight into how waste products are being cleared from CSF, characterizing CSF dynamics is crucial in advancing our knowledge of the brain’s clearance mechanisms.

### Limitations

The method developed in this work was optimized to study CSF dynamics in higher regions of the SAS, as these areas are traditionally thought to be the primary routes of CSF flow. Measuring CSF flow in other regions like the lateral or the fourth ventricle will be challenging due to flow-induced signal loss which may be inevitable since the method is aimed to study very subtle motions. A total of 6 healthy individuals were scanned all during the morning between 8 am and 12 am. It has been shown that CSF excretion varies over the circadian cycle[48], with the excretion being lowest in the morning. However, since the PC-MRI measurements performed in the aqueduct showed a non-zero net flow this would not explain the absence of net flow in the SAS. Although LFOs are currently not included in the underlying model of CSF dynamics for in vivo measurements, primarily due to the lack of an accurate recording device, the simulations indicated that the influence of LFOs would not affect the estimation of the net velocity (it would likely result in an underestimation of the other physiological processes) which also does not explain the absence of net flow.

## Conclusion

This study proposed and validated a novel method that is capable of measuring ultraslow net CSF velocity in the presence of much larger confounding factors from head motion and physiological pulsations. Measurements in the SAS revealed no detectable net flow of CSF towards the SSS. These findings challenge the classical view of CSF excretion and absorption and underscore the increasingly need for re-evaluation of traditional models of CSF dynamics to help advance the knowledge on brain clearance.

## Notes

### Competing Interest Statement

The authors have declared no competing interest.

