## Supplementary files for "Challenging the classical view of CSF flow: measuring CSF net velocity in the human subarachnoid space with 7T MRI"

**Supplementary info**

Supplementary Figure 1

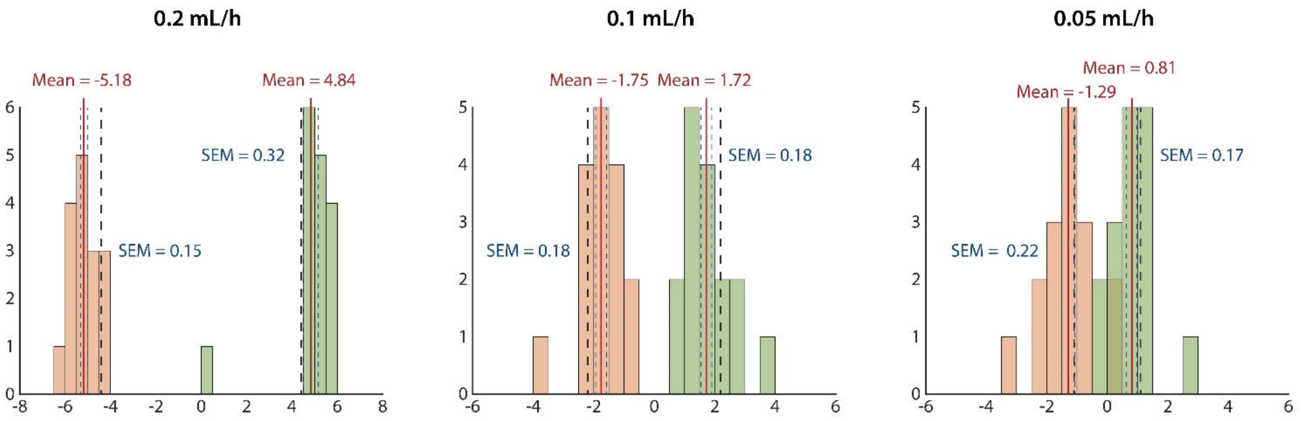


Histograms of the mean velocity of all slices of the motorized phantom for the different infusion rates: 0.5 mL/h (left), 0.1 mL/h (middle) and 0.05 mL/h (right). The red line indicates the mean velocity, the dashed blue line the SEM over all slices and the black dashed line the ground truth velocity.

Supplementary Figure 2

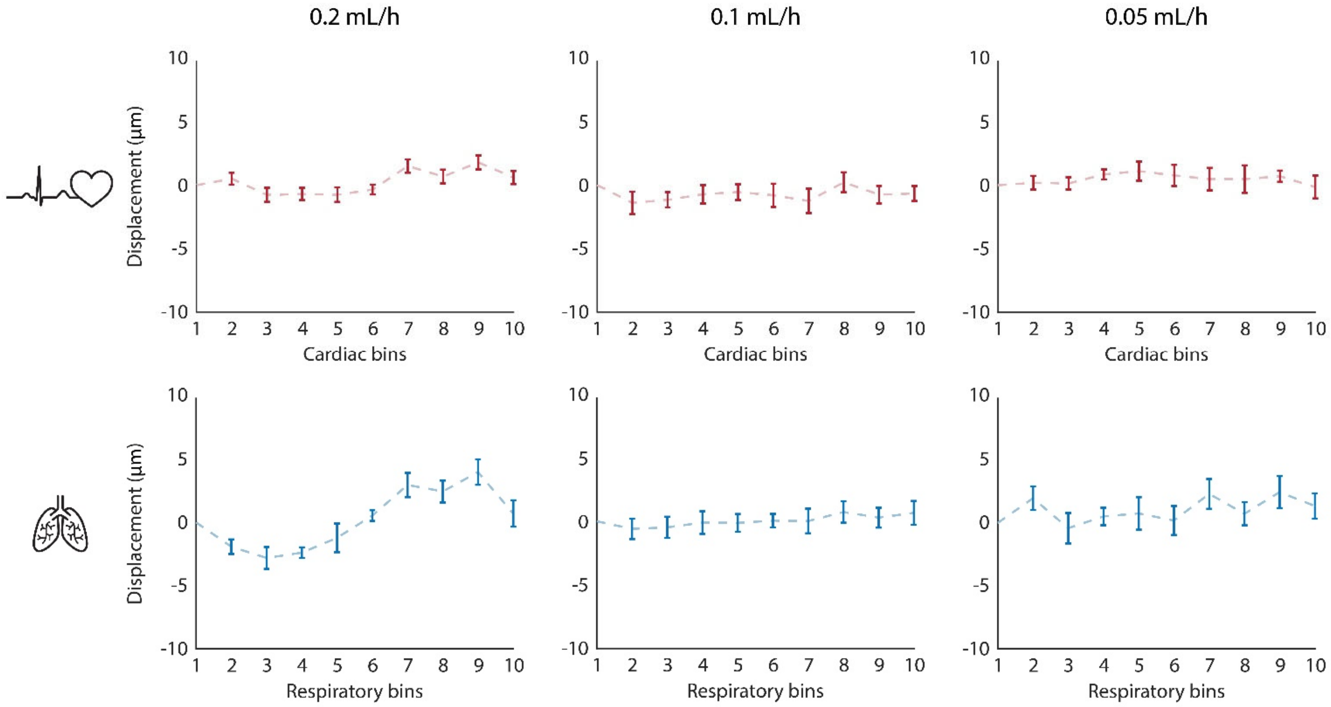


Results of voxelwise fit of the motorized flow phantom after motion correction and eddy current correction. The top row shows the cardiac motion and the bottom row the respiratory motion for different infusion rates: 0.2 mL/h (first column), 0.1 mL/h (middle column) and 0.05 mL/h (last column). The error bars indicate the SEM over all (n = 16) slices. The first bin is used as a reference point which has zero displacement and no SD by definition. Note the limits of the y-axis which are 10 or 20 times smaller than the residual fitted motion curves (Figure 6).

Supplementary Figure 3

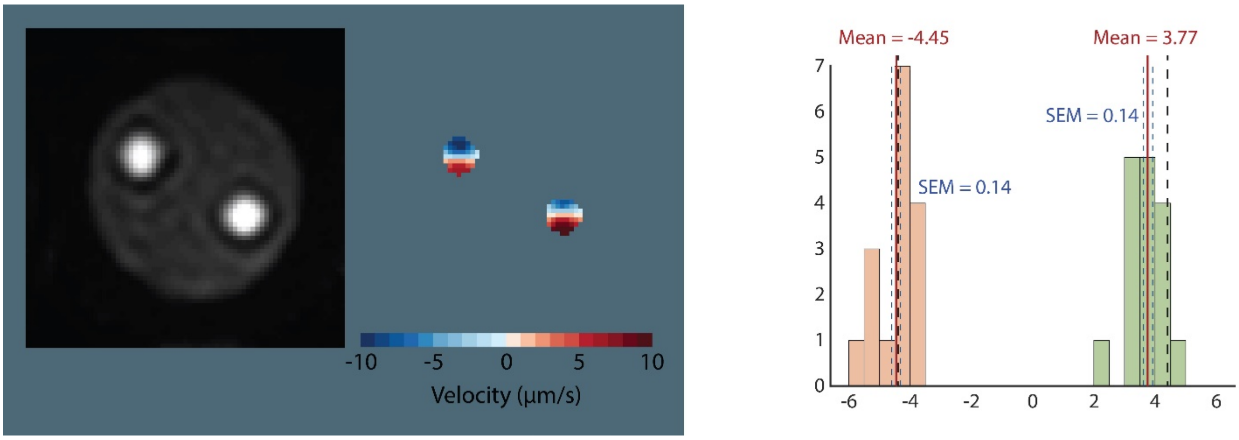


Results of the isolated phantom experiments placed at an incline. The net velocity maps show a similar gradient as was observed in the in vivo phantom experiments. The histograms shows the average net velocity in both tubes over all (n = 16) slices. The red line indicates the mean velocity, the dashed blue line the SEM over all slices and the black dashed line the ground truth velocity.

Supplementary Figure 4

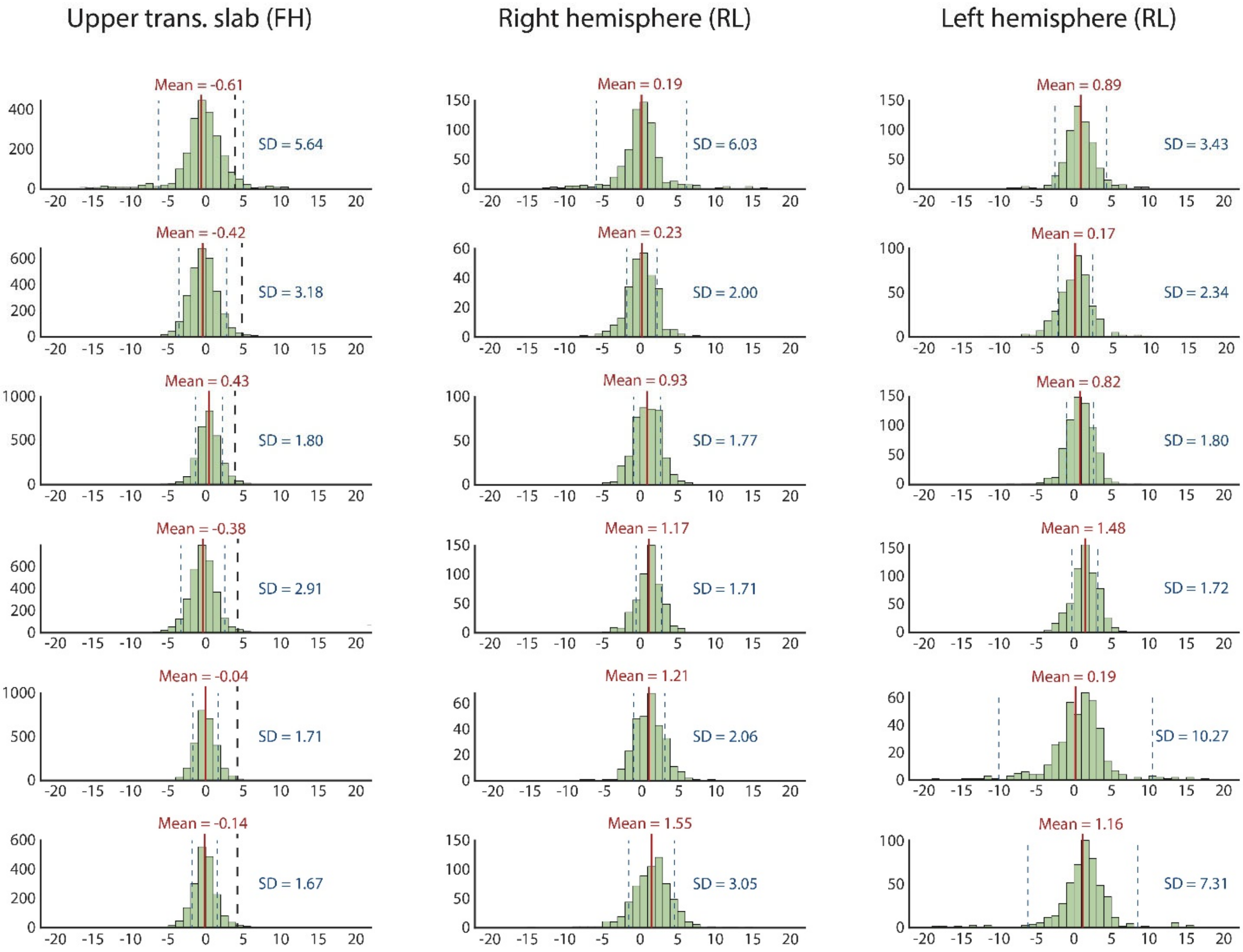


Histograms of the net velocity as estimated in 10 transverse slices of the SAS located just above the ears for FH encoding (first column) or 5 upper transverse slices of the SAS in the left and right hemisphere for the RL encoding (second and third column, respectively). The red line indicates the mean velocity and the blue dashed line the standard deviation (SD) estimated over all voxels within the ROI. The black dashed in the first column indicates the expected net velocity in the SAS based on CSF flow measured in the aqueduct. For visualization purposes, the histograms were cut off at [-20 20]. No clear net velocity towards the sagittal sinus was found, neither for FH nor for RL encoding.

Supplementary Figure 5

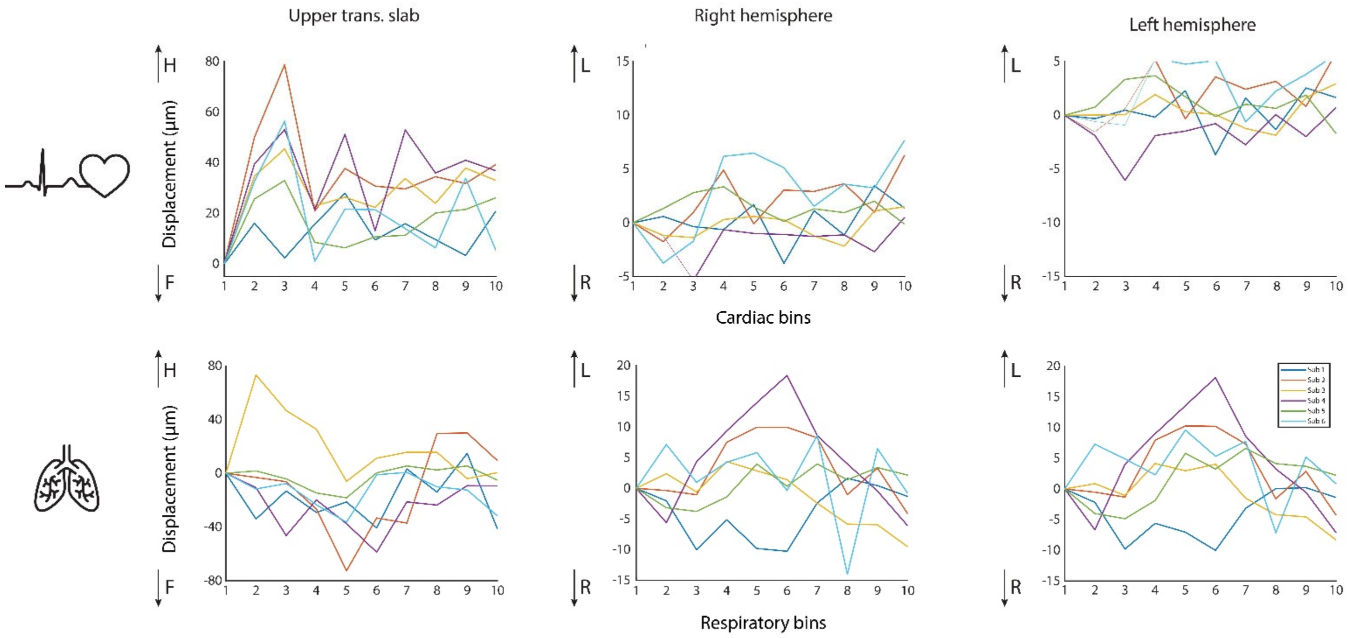


Estimated cardiac (upper row) and respiratory (bottom row) motion from voxelwise fit on motion correction residuals. The FH encoded motion is analyzed in the 5 upper transverse slices of the SAS (first column) and the RL encoded motion is analyzed in 10 transverse slices just above the ears in the right and left hemisphere (second and third column, respectively). Motion is binned into 10 bins which represent 5%, 15%, 25%, ... up to 95% of the cardiac or respiratory cycle and bin 1-5 represent inhalation and bin 6-10 exhalation. The first bin is used as a reference point which has zero displacement and no SD by definition.

Supplementary Table 1. Measured net CSF velocities in the SAS.

| | **Measured rightward net velocity** | **Measured leftward net velocity** |
|---|---|---|
| **Sub 1** | 0.9 | 0.2 |
| **Sub 2** | 0.2 | 0.2 |
| **Sub 3** | 0.8 | 0.9 |
| **Sub 4** | 1.5 | 1.2 |
| **Sub 5** | 0.2 | 1.2 |
| **Sub 6** | 1.2 | 1.6 |
| **Mean ± SEM** | 0.80 ± 0.21 | 0.88 ± 0.23 |
